# Mixed-stock analyses among migratory, non-native Chinook salmon at-sea and assignment to natal sites in freshwater at their introduced range in South America

**DOI:** 10.1101/732248

**Authors:** Selim S. Musleh, Lisa W. Seeb, James E. Seeb, Billy Ernst, Sergio Neira, Chris Harrod, Daniel Gomez-Uchida

**Affiliations:** Programa de Doctorado en Ciencias con Mención en Manejo de Recursos Acuáticos Renovables (MaReA), Faculty of Natural Sciences and Oceanography, P.O. Box 160-C, Concepción, Chile; Núcleo Milenio INVASAL, Concepción, Chile; School of Aquatic and Fishery Sciences, University of Washington, 1122 NE Boat Street, Box 355020, Seattle, WA 98195-5020, USA; Department of Oceanography, Universidad de Concepción, Faculty of Natural Sciences and Oceanography, P.O. Box 160- C, Concepción, Chile; Instituto de Ciencias Naturales Alexander von Humboldt, Facultad de Ciencias del Mar y Recursos Biológicos, Universidad de Antofagasta, Antofagasta, Chile; Universidad de Antofagasta Stable Isotope Facility, Instituto Antofagasta, Universidad de Antofagasta, Antofagasta, Chile; Genomics in Ecology, Evolution and Conservation Laboratory, Department of Zoology, Universidad de Concepción, Faculty of Natural Sciences and Oceanography, P.O. Box 160- C, Concepción, Chile

**Keywords:** naturalized Chinook salmon, mixed-stock analysis, invasion biology, fishery management of non-native species, northern Patagonia

## Abstract

Invasive species with migratory behavior and complex life cycle represent a challenge for evaluating natal sites among individuals. Private and government-sponsored initiatives resulted in the successful introduction and naturalization of Chinook salmon (*Oncorhynchus tshawytscha*) throughout northern and southern Patagonia in South America. These migratory fish breed in freshwater, but spend most of their life at sea feeding, forming abundant populations in several watersheds draining into the southeast Pacific Ocean. We used single nucleotide polymorphisms (SNPs) combined with genetic structure and mixed-stock analyses to evaluate natal sites of Chinook salmon at-sea caught in one estuary and two coastal locations compared to reference populations from breeding sites in freshwater. Firstly, Bayesian individual-assignment analyses revealed no genetic structure among adults caught off the coast of the Toltén River and migrating (maturing) adults caught in Toltén River estuary, suggesting they likely belong to a single population. Secondly, mixed-stock genetic analyses revealed that most at-sea Chinook salmon caught in one estuary and two coastal locations likely originated from spawners from the nearest river (90-95%), with a small contribution from adjacent watersheds (5-10%). This appears consistent with Chinook salmon populations in their native range in which juveniles migrate short distances (100-200 km) from their river of origin to coastal feeding grounds, some of which became donor of propagules for non-native Chinook salmon populations under study. Mixed-stock genetic analyses provide considerable potential to identify the population of origin of Chinook salmon mixtures caught off the coast. They also seem an appropriate proof of concept to help identify potential immigrants from other watersheds as well as migration patterns and invasion pathways in a non-native species.

## Introduction

Population genetics and genomics approaches have been increasingly implemented for studying invasions and their ecological and evolutionary consequences (Barrett 2015; Crommenacker et al. 2015). Primary foci among studies include comparing genetic metrics between native and non-native populations, identifying invasion pathways from donor population sources, and evaluating the role of receiving environments on selection and adaptation among established genotypes (Barrett 2015; Chown et al. 2015). Applications oriented to assign individuals of unknown origin to their natal locations in their non-native, post-establishment range have received lesser attention (but see Darling and Folino-Rorem 2009 for an exception). Knowledge of migration patterns among invasive species with a complex life cycle might benefit from such applications, namely species in which juveniles and adults occupy different environments. Also, and because adults may form population mixtures, quantifying proportions from different populations is a crucial first step to gauge ecological impacts of specific non-native populations, as prey or predators, on native ecosystems, and to recommend potential management actions.

Chinook salmon (*Oncorhynchus tshawytscha*) is an anadromous fish native to the Northern Hemisphere and widely distributed in the North Pacific Ocean. Chinook salmon breed in freshwater but feed most of their lives at sea. In their native range, Chinook salmon juveniles migrate to the ocean either in the spring or summer after emerging from the gravel (subyearling) or outmigrate the next spring (yearling), typically followed by 2 to 5 years growing at sea - (Quinn 2005). Patterns of Chinook salmon migration are a highly variable trait among populations; some undertake long migrations, while others remain close to the coast (Healey 1991; Trudel et al. 2009). Chinook salmon are carnivorous fish that prey over a broad range of pelagic species such as euphausiids, decapods, cephalopods and fishes (Brodeur 1990; Daly et al. 2009; Riddell et al. 2018). The species is also an important resource for recreational, tribal, and commercial fisheries and is considered one of the most valuable Pacific salmon species in North America (Quinn 2005).

The clear fishery potential of Chinook salmon triggered government-sponsored and private initiatives to successfully propagate it throughout Pacific Ocean watersheds in South America during 1978-1990 (Correa, Gross 2008), with donor (native) populations spanning multiple geographic regions from North America (Correa, Moran 2017; Gomez-Uchida et al. 2018; Riva Rossi et al. 2012). A recent genetics study has shown that non-native Chinook salmon have invaded new watersheds in South America establishing genetically differentiated population groups originating from both artificial and natural dispersal (Gomez-Uchida et al. 2018). Genetically isolated populations can be useful as reference groups to understand migratory behavior and identifying natal populations among mixed stock salmonids at sea (Bellinger et al. 2015; Bradbury et al. 2016; McKinney et al. 2017). Similar applications could be implemented in their introduced range for understanding key aspects of invasion ecology, namely pathways, sources of propagules, and the relative contribution of specific populations.

Here we employed single nucleotide polymorphisms (SNPs) and individual assignment methods to characterize genetic structure and population mixtures among non-native Chinook salmon at sea, including one estuary and two coastal locations. We probabilistically assign at-sea individual genotypes to natal sites in freshwater using a set of reference populations. Our study design includes two locations in South America: one study site off Toltén River watershed (39°S) and another at Gulf of Ancud (42°S) in northern Patagonia (Figure 1). These represent two potentially important feeding grounds for non-native, migratory salmonids. Our goal was to help understanding ecology of non-native Chinook salmon, especially in relation to the marine phase and ocean migration patterns, which remain largely unstudied.

**Figure 1.**
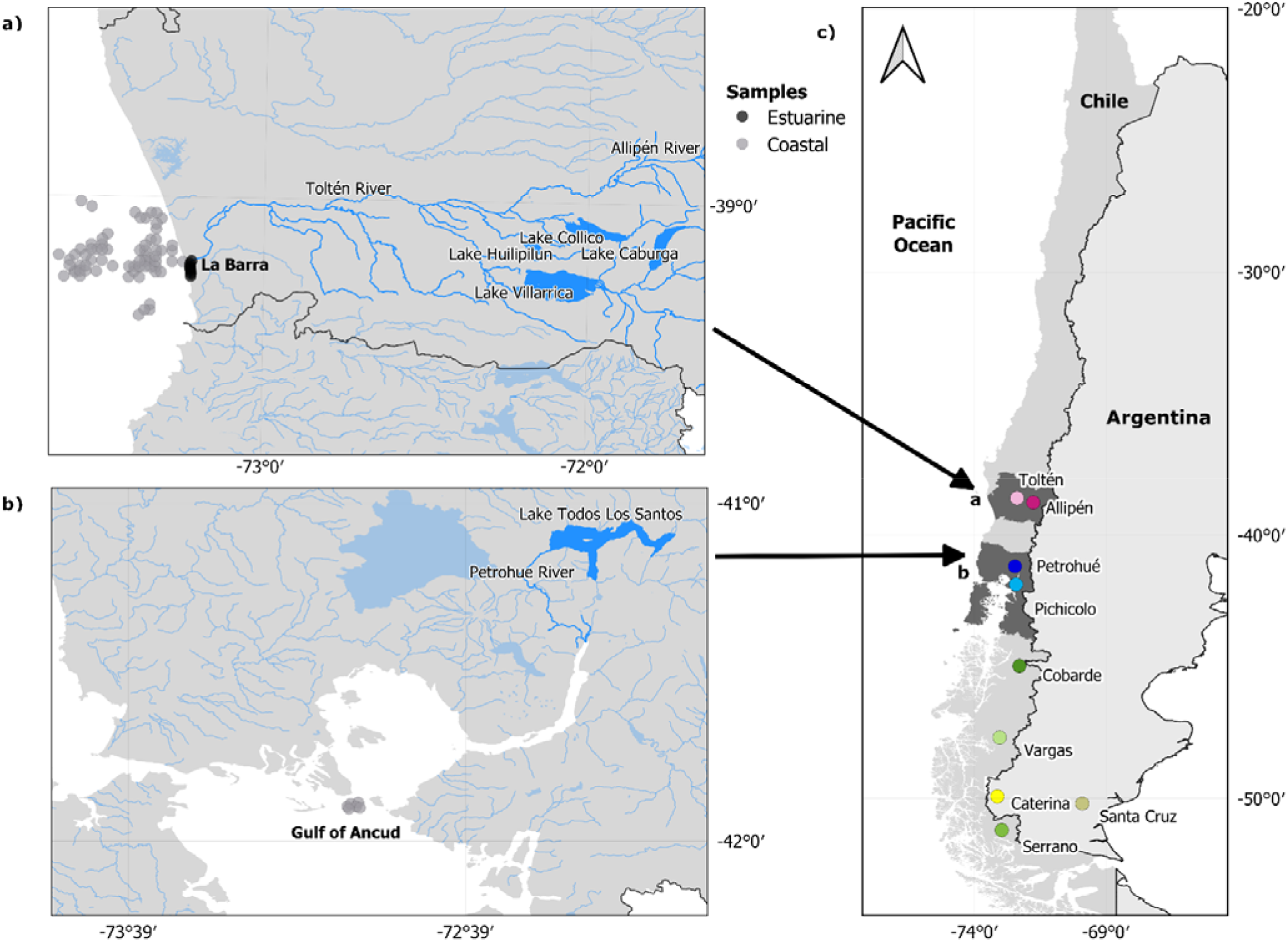
Current and historical collection sites of non-native Chinook salmon in South America. **a.** At-sea sampling sites from Araucanía Region. Grey dots are coastal fishing sites; black dots are fishing sites at the estuary of Toltén River. **b.** At-sea sampling sites from Los Lagos Region. Grey dots are fishing points at Gulf of Ancud. **c.** Natal freshwater sites in Pacific Ocean and Atlantic Ocean watersheds.

## Materials and Methods

### Study sites and sampling

Toltén River supports one of the largest Chinook salmon populations identified to-date in Chile (Gomez-Uchida et al. 2018). The entire watershed comprises four lakes and multiple rivers that run from the Andean mountains to the Pacific Ocean (Figure 1a). River habitats in the western foothills of the Andes (1500 – 2000 masl) provide high-quality habitat for breeding (Gomez-Uchida et al. 2016). The coastal area off Toltén River is highly productive and supports large stocks of common sardine (*Strangomera bentincki*) and anchovy (*Engraulis ringens*) (Instituto de Fomento Pesquero 2018), an important prey item for coastal Chinook salmon (Gomez-Uchida et al. 2016). Chinook salmon samples from the estuary (n=121) and off the coast of Toltén River (n=53) were collected by small-scale artisanal fishers under Chile’s Undersecretariat of Fisheries & Aquaculture permit (Res. Ex. 3417/2014) during austral summer 2015 (Table 1).

**Table 1.** Location, date, sample size (*n*), and genetic statistics (expected heterozygosity, ***H_E_***; allelic richness, ***A_R_***) for non-native Chinook salmon

| Location | Date | $n$ | $H_E$ | $A_R$ |
| --- | --- | --- | --- | --- |
| **Toltén River, estuary** | December 2014 - January 2015 | 121 | $0.279 \pm 0.013$ | **$1.895 \pm 0.022$** |
| **Toltén River, coastal** | January 2015 | 53 | $0.272 \pm 0.014$ | **$1.785 \pm 0.014$** |
| **Gulf of Ancud, coastal** | January 2014 | 42 | $0.293 \pm 0.012$ | **$1.932 \pm 0.012$** |

The Gulf of Ancud is located within the commonly referred to Inner Sea of Chiloé (FIGURES Figure 1b) in Chile’s northern Patagonia (Los Lagos Region). This marine area consists of four micro-basins interconnected through narrow passes between islands. Gulf of Ancud has been described as a breeding and nursery area for several native fishes (Bustos et al. 2008) and is therefore a potential feeding ground for juvenile and adult Chinook salmon. Samples (n=42) originated from bycatch of the southern hake (*Merluccius australis*) long-line fishery taken during 2014 austral summer (Table 1).

### SNP genotyping and selection; genetic statistics

Genomic DNA was isolated from each Chinook salmon axillary process (i.e., a small pointed protrusion located above the pelvic fin) using a Macherey-Neigel Nucleospin Tissue kit following manufacturer’s instructions. PCR was carried out using Fluidigm^®^ 96.96 dynamic array chips following established pre-amplification and amplification protocols (Seeb et al. 2009; Smith et al. 2011). We screened DNA samples through a suite of 170 SNPs (Table S1). These loci were selected based on information content among non-native populations, fit to Hardy-Weinberg equilibrium (HWE) proportions, and non-significant linkage disequilibrium (LD) among loci (Gomez-Uchida et al. 2018). We calculated genetic statistics, expected heterozygosities and allelic richness, for all three groups of samples using GENEPOP (Rousset 2008).

### Individual-based inference of genetic structure among Chinook salmon at Totten River

We assessed genetic population structure among estuarine and coastal Chinook salmon at Toltén River using Bayesian clustering based on minimization of HWE-LD departures implemented in STRUCTURE (Pritchard et al. 2000). This was based on demographic differences between these zones; previous surveys (Gomez-Uchida et al. 2016) indicated that coastal adults, especially females, were younger (ages 2+ and 3+) than estuary adults (ages 3+ and 4+). We therefore evaluated the likelihood that adults from estuary and coastal fisheries belong to one, two, or three different demes (number of demes = K values). For each K value, 10 independent iterations were run, using the admixture model with 20,000 iterations burn-in period of and 250,000 Markov Chain Monte Carlo steps after the burn-in. We arrived to “consensus” coefficients of individual membership (Q-values) within K values using CLUMMP to avoid label switching artifacts and multimodality among independent runs (Jakobsson, Rosenberg 2007).

### Mixed stock analyses of estuarine and coastal Chinook salmon and their assignment to reference populations in freshwater

We built a set of reference populations from Chinook salmon genotypes (n=504) including six Pacific Ocean and two Atlantic Ocean watershed sites in South America (Gomez-Uchida et al. 2018; Figure 1c). Multilocus genotypes at 170 SNPs (Table S1) originated from collections of juvenile and spawners taken from natal sites in freshwater. First, we performed a discriminant analysis of principal components (Jombart et al. 2010) to define the most probable number of reference populations. Second, we performed leave-one-out simulations to estimate the probability of assignment of genotypes from their natal sites to reference populations (i.e., self-assignment) using ONCOR (Kalinowski et al. 2008). Genotypes were bootstrapped by resampling alleles following Anderson et al. (2008) in 200 mixtures per natal site containing 100 individuals each. Highly admixed genotypes (possibly hybrids; Gomez-Uchida et al. 2018) were removed from the database, because in simulations they assigned to two or more reference populations. Third, we used Bayesian assignment (Rannala, Mountain 1997) to estimate membership probabilities of one estuary and two coastal Chinook salmon mixtures to reference populations using ONCOR.

## Results

### SNP genotyping and selection; genetic statistics

Expected heterozygosities among Chinook salmon from one estuary and two coastal locations varied between 0.272 and 0.293, while allelic richness varied between 1.785 and 1.932 (Table 1).

### Individual-based inference of genetic structure among Totten River Chinook salmon

Bayesian clustering analysis showed no evidence of genetic structure among migrating Chinook salmon adults caught at the estuary of Toltén River and adults caught off the coast. Individual membership coefficients were symmetric assuming *K*= 2 and *K*= 3 number of demes (Figure 2), consistent with the hypothesis that estuarine and coastal Chinook salmon belong to a single population off Tolten River.

**Figure 2.**
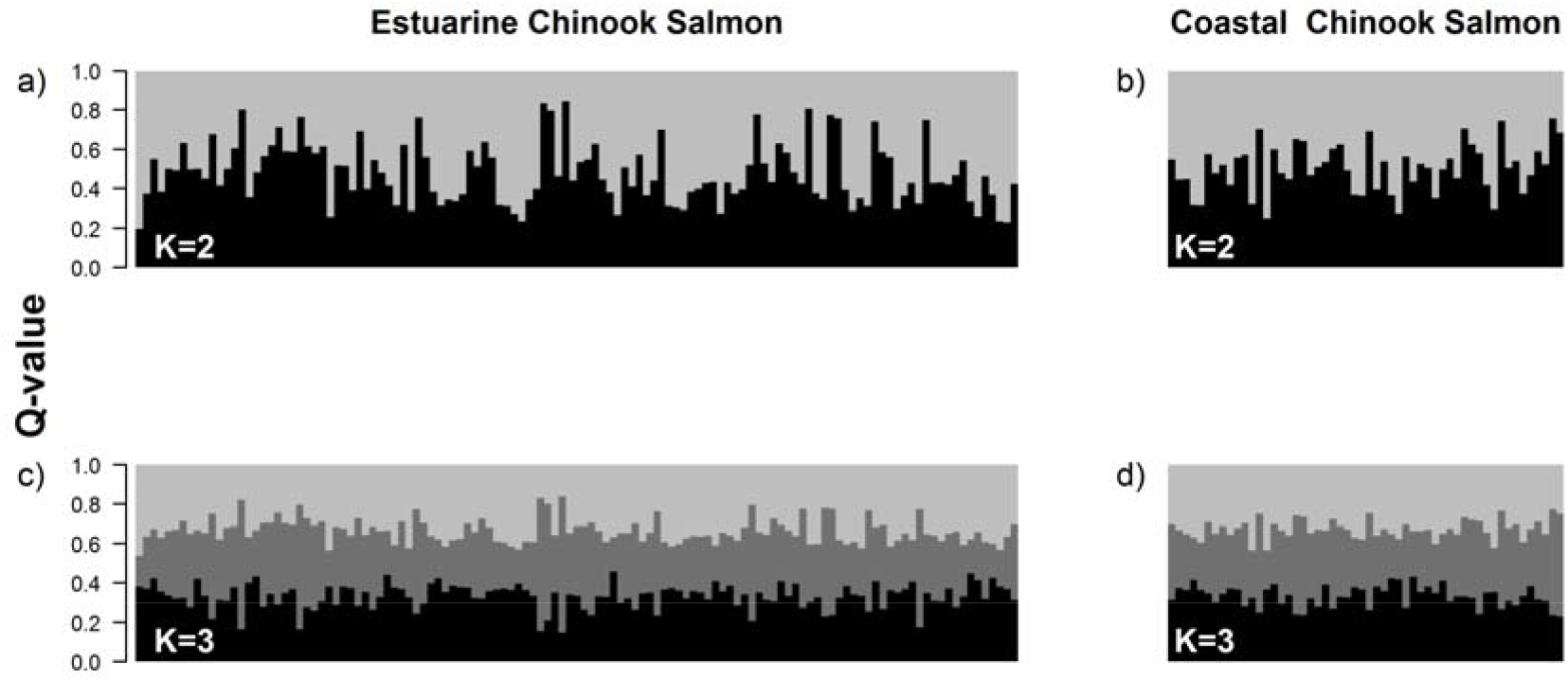
Individual ancestry coefficients (Q-values) among Chinook salmon genotypes sampled at the estuary (**a** and **c**) and off the coast of Tolten River (**b** and **d).** Clustering was performed assuming *K*=2 (**a** and **b:** upper panels) or *K*=3 demes (**c** and **d:** lower panels).

### Mixed stock analyses of estuarine and coastal Chinook salmon and their assignment to reference populations in freshwater

Discriminant analysis of principal components conducted on the dataset defined four genetically differentiated clusters, three among Pacific Ocean watersheds and one among Atlantic Ocean watersheds: (1) *Araucanía*, (2) *Los Lagos*, (3) *Aysén*, and (4) *Santa Cruz* (Figure 3a). We considered these clusters our reference populations, which were used for subsequent analyses. Leave-one-out simulations provided self-assignment values of 100% for most natal sites among Pacific Ocean watersheds; exceptions were simulations among Atlantic Ocean sites that self-assigned between 82% and 93% (Table 2).

**Figure 3.**
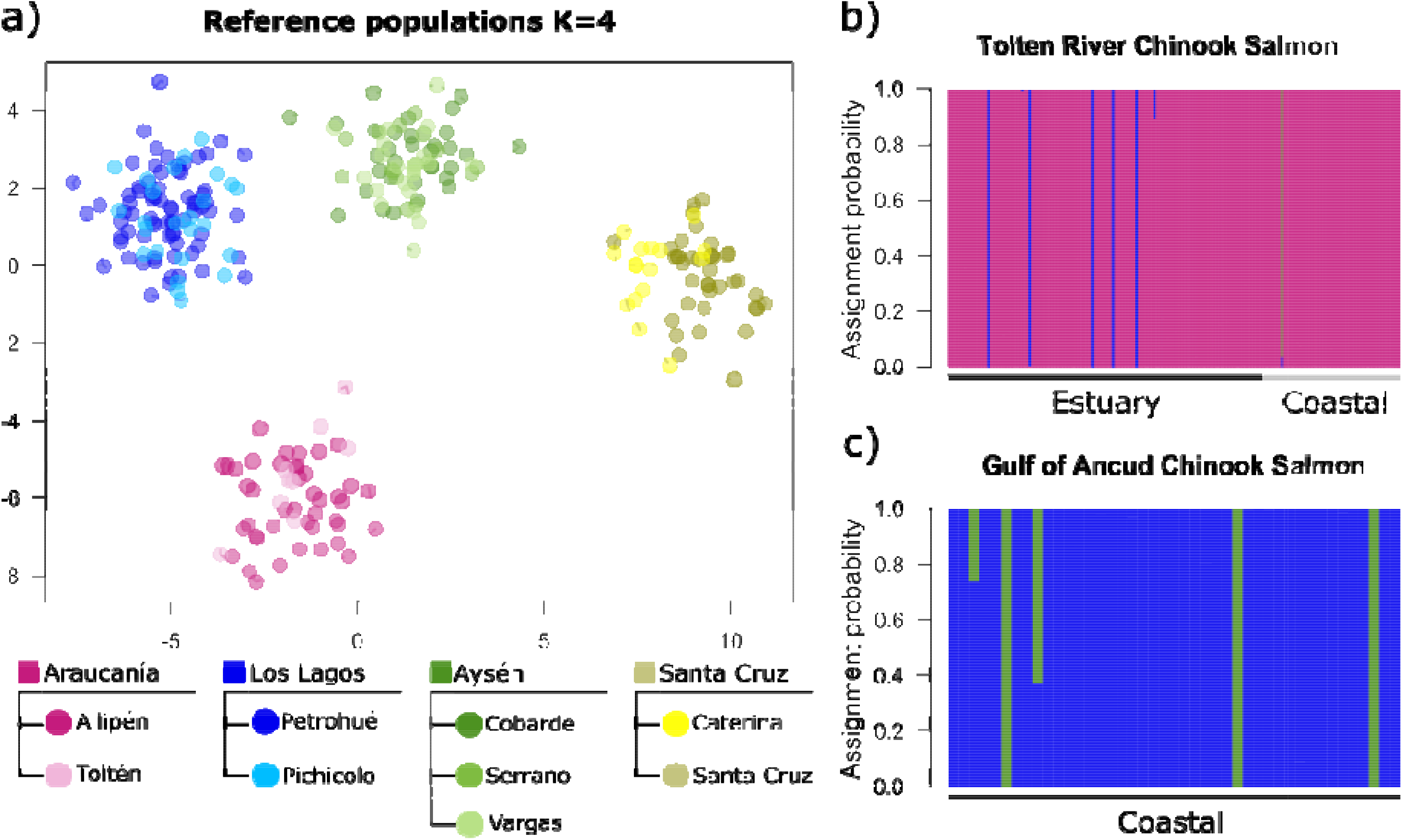
Reference populations in South America and assignment of Chinook salmon from two coastal and one estuary locations, **a)** Dots represent individual genotypes and dashed lines delimit reference populations. Colors represent natal sites (Table 2); squares are reference populations composed of two or more natal sites. **b)** Individual assignment probability of estuarine and coastal Chinook salmon off Toltén River to reference populations. **c)** Individual assignment probability of Chinook salmo from Gulf of Ancud to reference populations. Each bar represents an individual and colors represent probabilities to be assigned to one of four reference populations.

**Table 2.** Set of reference populations of Chinook salmon from South America based on Gomez-Uchida et al. (2018a). Colored dots represent the natal sites in freshwater; these are nested within four colored squares depicting genetically distinct reference populations. Self-assignment depicts the probability of an individual genotype from a specific natal site to be assigned to its reference population in simulations.

| *Basin* | *Country, Region* | *Natal sites* | *Reference population* | *Self-assignment* |
| --- | --- | --- | --- | --- |
| *Pacific Ocean* | Chile, Araucanía | ● Allipén | ■ Araucania | 100% |
|  | Chile, Araucanía | ● Toltén |  | 100% |
|  | Chile, Los Lagos | ● Petrohué | ■ Los Lagos | 100% |
|  | Chile, Los Lagos | ● Pichicolo |  | 100% |
|  | Chile, Aysén | ● Cobarde | ■ Aysén | 100% |
|  | Chile, Aysén | ● Vargas |  | 100% |
|  | Chile, Magallanes | ● Serrano |  | 100% |
| *Atlantic Ocean* | Argentina, Santa Cruz | ● Caterina | ■ Santa Cruz | 93% |
|  | Argentina, Santa Cruz | ● Santa Cruz |  | 82% |

The greatest proportion of Chinook salmon mixtures at-sea, from one estuary and two coastal locations, was assigned to the nearest reference population or rivers. Most estuarine and coastal individuals (>95%) caught off Tolten River were assigned to *Araucanía* population; five individuals caught at the estuary could be reliably assigned to *Los Lagos* population, whereas one individual caught off the coast could be assigned to *Aysén* population (Figure 3b). We obtained a similar result for Chinook salmon captured at Gulf of Ancud, with most genotypes (>90%) assigned to *Los Lagos* population and few individuals with genetic background from *Aysén* population (Figure 3c).

## Discussion

Using a combination of statistical methods to analyze population mixtures and genetic information provided by SNPs, we probabilistically assigned non-native Chinook salmon caught at sea to some of their natal sites in freshwater in South America. Mixed-stock analyses of samples from one estuary and two coastal locations suggested that most adults originated from the nearest river (90-95%), with limited contribution from adjacent watersheds (5-10%). Contributions from adjacent river sites likely represent immigrants. Below we discuss implications of our findings to illuminate ocean ecology and migration patterns among non-native Chinook salmon. Identifying such patterns has the potential to clarify invasion pathways as well as assist management of invasive species as we argue in the next paragraphs.

### Individual-based inference of genetic structure among Tolten River Chinook salmon

We used clustering methods to examine putative genetic structure among Chinook salmon captured in the estuarine and coastal zones off Toltén River, but no genetic structure was apparent amid demographic differences between these zones. This suggests that coastal and estuarine adults are composed of different age classes and likely belong to one single, self-sustaining population. We speculate that Tolten River Chinook salmon has been established for four to five generations since migrating adults showed in high numbers during 2000-2005, assuming a generation length of four years (Gomez-Uchida et al. 2016). Time since establishment may thus be insufficient to evolve neutral genetic structure for a single, large watershed. Suk et al. (2012) reported evolution of genetic structure within 10 generations among non-native Chinook salmon that invaded Lake Huron’s watershed in North America, while Kinnison et al. (2002) found genetic differentiation within 30 generations among introduced Chinook salmon to New Zealand, which originated from a single donor region. Yet, and albeit beyond to the scope of this study, Tolten River Chinook salmon may continue to experience selection at adaptive loci in response to environmental selection imposed by a novel environment. Adaptation following invasion may be linked to changes in phenotypic traits as demonstrated in other invasive taxa, including fishes (Monzon-Arguello et al. 2014), mammals (White et al. 2013) and plants (Vandepitte et al. 2014).

### Mixed stock analyses of estuarine and coastal Chinook salmon and their assignment to reference populations in freshwater

We demonstrated that most Chinook salmon adults sampled from one estuary and two coastal sites originated from the nearest reference population and natal sites in freshwater. Clear differences among four reference populations enable individual assignment of adults captured at sea to their natal sites in freshwater. This resulted from genetic divergence among donor Chinook salmon lineages from North America that is still trackable, combined with contemporary changes driven by founder effects, genetic drift, and gene flow, forming genetically isolated populations (Gomez-Uchida et al. 2018). Although self-assignment was high, it should be noted that our coverage of all potential natal sites in freshwater was incomplete. For example, we had no Chinook salmon genotypes from natal sites at Valdivia and Bueno rivers, two watersheds located south of Tolten River, but north of Petrohué River (Fig. 1c). However, we hypothesize that colonization of Valdivia and Bueno watersheds by Chinook salmon possibly resulted from natural dispersal from Toltén River, Petrohue River, or both. This implies genetic differentiation between sampled and unsampled populations may be weak and likely to belong to the same reference population (e.g., Bradbury et al. 2016).

What is the evidence supporting patterns of coastal ocean migration among Chinook salmon? Large-scale ocean surveys of tagged juveniles in their native range indicate that most will remain within 100-200 km of their natal river in their first year, even though some may travel thousands of km north during their second year, depending on life history variation among populations (Trudel et al. 2009). Tagging studies and mixed-stock genetic analyses among native Chinook salmon adults tell a similar story – a great proportion were recovered or caught within major regions where they were tagged or assigned to, although with notable exceptions (Bellinger et al. 2015; Weitkamp 2010). We thus predict that an important proportion of non-native Chinook salmon adults may remain near their natal watershed feeding on the continental shelf; additionally, other adults may experience long migrations along the Pacific Ocean coast off Chile, and possibly offshore, but return to their natal watersheds before their breeding season. This is consistent with several lines of evidence. First, abundance of suitable prey items such as sardine and anchovies may represent the main factor driving Chinook salmon to remain in coastal habitats close to their river of origin in Chile, especially off Tolten River, where sardine abundance is one of the highest off Chile’s coast (Bustos et al. 2008; Gomez-Uchida et al. 2016; Instituto de Fomento Pesquero 2018). Second, Chinook salmon from Tolten River (reference population: *Araucanía*) and Petrohue River (reference population: *Los Lagos*) were likely founded from donor (native) populations that experience short migration routes during juvenile stages after Trudel et al. (2009), namely coastal Oregon and Puget Sound populations, respectively (Gomez-Uchida et al. 2018). Third, Chinook salmon adults have been caught by recreational fishers in both Pacific Ocean and Atlantic Ocean, in some cases thousands of km away from the nearest established population in freshwater, suggesting ‘stray’ salmon may migrate long distances (Liotta 2019; authors’ field observations).

Invasive species frequently experience range expansions and distributional changes, and predicting invasion pathways as well as evaluating the rate of spread are crucial for efficient management of non-native taxa (Sakai et al. 2001). Mixed stock analyses hold clear potential to study migration patterns among non-native Chinook salmon by assigning adults at-sea to their natal sites in freshwater. Such analyses may clarify which non-native populations may be acting as source of propagules, particularly long-distance migrants as part of an ongoing process of invasion by this successful salmonid.

## Acknowledgements

We are indebted to Cristian Canales-Aguirre, Pablo Rivara, Diego Cañas, Mauricio Cañas, and Francisca Valenzuela-Aguayo for their substantial contribution to sample collection. Carita Pascal performed all laboratory procedures, including SNP genotyping, allele scoring, and database management. This research was supported by Núcleo Milenio INVASAL funded by Chile’s government program, Iniciativa Cientifica Milenio from Ministerio de Economia, Fomento y Turismo, and Chile’s government grants, FONDECYT 1130807, 1191256, and FIPA 2014-87. Student support came from CONICYT 21160640 doctoral scholarship to SSM.

